# Real-Time Observation of Structure and Dynamics during the Liquid-to-Solid Transition of FUS LC

**DOI:** 10.1101/2020.10.19.345710

**Authors:** Raymond F. Berkeley, Maryam Kashefi, Galia T. Debelouchina

## Abstract

Many of the proteins found in pathological protein fibrils also exhibit tendencies for liquid-liquid phase separation (LLPS) both *in vitro* and in cells. The mechanisms underlying the connection between these phase transitions have been challenging to study due to the heterogeneous and dynamic nature of the states formed during the maturation of LLPS protein droplets into gels and solid aggregates. Here, we interrogate the liquid-to-solid transition of the low complexity domain of the RNA binding protein FUS (FUS LC), which has been shown to adopt LLPS, gel-like, and amyloid states. We employ magic-angle spinning (MAS) NMR spectroscopy which has allowed us to follow these transitions in real time and with residue specific resolution. We observe the development of β-sheet structure through the maturation process and show that the final state of FUS LC fibrils produced through LLPS is distinct from that grown from fibrillar seeds. We also apply our methodology to FUS LC G156E, a clinically relevant FUS mutant that exhibits accelerated fibrillization rates. We observe significant changes in dynamics during the transformation of the FUS LC G156E construct and begin to unravel the sequence specific contributions to this phenomenon with computational studies of the phase separated state of FUS LC and FUS LC G156E.

**Significance:** The presence of protein aggregates and plaques in the brain is a common pathological sign of neurodegenerative disease. Recent work has revealed that many of the proteins found in these aggregates can also form liquid-liquid droplets and gels. While the interconversion from one state to another can have vast implications for cell function and disease, the molecular mechanisms that underlie these processes are not well understood. Here, we combine MAS NMR spectroscopy with other biophysical and computational tools to follow the transitions of the stress response protein FUS. This approach has allowed us to observe real-time changes in structure and dynamics as the protein undergoes these transitions, and to reveal the intricate effects of disease-relevant mutations on the transformation process.

## Introduction

Neuronal protein aggregates are hallmarks of neurodegenerative disease whose biological genesis is still poorly understood^1,2^. While their cellular functions may vary, proteins found in these aggregates often contain intrinsically disordered domains with low complexity charge-patterned sequences. These sequences not only promote the adoption of fibrillar or aggregated states but are also important drivers for liquid-liquid phase separation (LLPS) and the formation of membraneless organelles^3,4^. As such, many proteins that are found in pathological neuronal aggregates can exist in a range of biophysical states that span a broad range of dynamic regimes including liquid-liquid droplets, hydrogels and amyloid fibrils. Some examples include proteins associated with Alzheimer’s disease (tau)^5–10^, Parkinson’s disease (α-synuclein)^11–14^, and frontotemporal lobar degeneration (TDP-43 and FUS)^15,16^.

Recent work has started to establish connections between these biophysical states *in vitro*. For example, the repeated coacervation and dissolution of liquid droplets of RNA-binding proteins such as hnRNPA1 has been shown to accelerate the formation of protein aggregates^17^. Imaging experiments often show fibrils that grow from the center of TDP-43, FUS and hnRNPA1 droplets suggesting that phase separated protein environments may serve as centers for the nucleation and growth of amyloid in the presence or absence of RNA^18–20^. Reversible amyloid-like fibers of FUS segments have also been found embedded within hydrogels^21^. Although it is known that phase separation is not a requirement for the growth of fibrils *in vitro*^6^, the growing body of evidence linking mature LLPS protein droplets and hydrogels to aggregates and fibrils suggests that the aberrant transition between these states could be a physiologically relevant factor in disease pathogenesis^7,16,19^.

The current biophysical toolbox contains well-established methodologies that can characterize each of those states separately. For example, liquid droplets have been analyzed with a variety of imaging, spectroscopic, and computational approaches that have provided valuable information regarding the sequence requirements and the nature of protein-protein interactions that drive LLPS^20,22–34^. On the other hand, magic angle spinning (MAS) NMR spectroscopy and cryo-EM have revealed the common structural principles that underlie the formation of stable, β-sheet rich amyloid fibrils^5,16,35–37^. While more challenging due to their viscous, dynamic and heterogeneous nature, hydrogels have also been amenable to characterization by MAS NMR approaches^38,39^. Yet, to establish a comprehensive view of how these states are connected on the molecular level and how these connections may break in disease, requires a strategy that can ideally observe these transformations in real time and in the same sample at atomic resolution. Here, we explore the capability of MAS NMR spectroscopy to achieve this goal.

MAS NMR spectroscopy is a versatile structural technique that allows the investigation of biological samples of different sizes, complexity and material state^40,41^. While such samples may experience line broadening interactions due to slow molecular tumbling, strong dipolar interactions and chemical shift anisotropy, the line broadening can be removed by spinning the sample at 54.7° (the magic angle)^42,43^. During magic angle spinning, various pulse sequences can reintroduce the magnetic spin interactions in a controlled manner that allows the extraction of structurally meaningful information such as the chemical environment and the atom identity, the protonation state, the distance between atoms and their relative orientation. Information can be reintroduced based on the scalar couplings between covalently bonded atoms, similar to solution NMR, enabling the description of the mobile components in the sample^44^. On the other hand, dipolar-based pulse sequences can be used to detect protein regions or sample components that experience slow motions and detectable dipolar interactions through space^45^. The combination of the two approaches provides the opportunity to dynamically edit the NMR spectra and to describe both the mobile and rigid components of the sample^46^.

In previous work, we have used MAS NMR spectroscopy to follow the transition from the liquid droplet to the gel state of the chromatin related protein HP1α^47^. This approach allowed us to detect specific serine residues that experience large changes in mobility and that appear to be important for the formation of cross-linking interactions in the gel state. Here, we extend this approach to the low complexity (LC) domain of the RNA-binding protein *Fused in sarcoma* (FUS) and we follow the transformation of individual samples of FUS LC from the liquid droplet to the gel and amyloid states in real time. We also compare the wild-type sequence and a sequence that harbors a pathogenic G156E mutation that greatly accelerates the onset of FUS fibrils *in vitro* and increases the likelihood for the development of ALS-spectrum diseases in patients^19^. We complement these studies with coarse-grained simulations to uncover the subtle changes in protein-protein interaction propensities that are amplified through this transformation process.

## Materials and Methods

### Expression of FUS LC and related constructs

FUS LC was prepared from recombinant Rosetta™ (DE3) competent *E. coli* cells (Novagen) that had been transformed with a plasmid (Addgene #98653) encoding the 6xHis-MBP-TEV-FUS(1-163) sequence^48^. Seed cultures were grown to saturation from freshly transformed colonies and inoculated at a 1% v/v ratio into either LB or ^15^N/^13^C-M9 medium. The cultures were grown at 37°C to an OD600 of ~ 0.7 and protein expression was induced by the addition of 1 mM isopropyl-β-D-thiogalactoside. The cultures were allowed to express protein for 4 hours at 37°C before being harvested by centrifugation at 10,000 RCF and 4°C for 30 minutes. After decanting the supernatant, cell pellets were stored at −80°C for later use. In addition to the wild-type FUS LC fusion protein, the pathogenic mutant FUS LC G156E protein was prepared. To generate the mutant, the requisite G to E mutation was introduced to the wild-type FUS LC plasmid using a NEBuilder^®^ HiFi DNA Assembly Cloning Kit. Expression and purification conditions of the G156E mutant protein were identical to those of the wild type.

### Purification of FUS LC and related constructs

Here, we followed the published protocol by Burke *et al.* with some modifications^31^. The frozen cell pellet was thawed, resuspended in lysis buffer (20 mM sodium phosphate, 300 mM sodium chloride, Roche cOmplete™ EDTA-free Protease Inhibitor Cocktail, pH 7.4, 4°C) and lysed by pulsed sonication for 30 minutes using a Qsonica sonicator with a 1/8” diameter probe tip at 12 kHz (60%) output and 4°C. The lysate was cleared by centrifugation at 20,000 RCF for 30 minutes and the supernatant was incubated with Thermo Scientific™ HisPur™ Ni-NTA Resin for 30 minutes at 4°C. The suspension of beads was washed with 10 column volumes of lysis buffer containing 10 mM imidazole, and protein was eluted with 2 column volumes of lysis buffer containing 250 mM imidazole. The eluted protein was incubated with 6xHis-tagged Tobacco Etch Virus (TEV) protease at a ratio of 1:150 TEV to MBP-FUS LC for 5 hours and 25°C in order to cleave MBP from FUS LC. FUS LC crashes out upon cleavage to yield a cloudy suspension with white clumps of aggregated protein. After TEV cleavage, 8 M urea was added to the reaction mixture to solubilize the aggregated FUS LC. The TEV reaction was monitored by SDS-PAGE (note that FUS LC does not bind Coomassie, so cleavage was verified by the gel shift of the MBP band). The mixture containing solubilized FUS LC was diluted to 2 mg/ml with 20 mM CAPS, 150 mM sodium chloride, pH 11 and subjected to size exclusion chromatography over a GE HiLoad^®^ 16/600 Superdex^®^ 75 pg column. This protocol generally yields 10-15 mg of FUS LC per liter of culture in both LB and M9 media.

### Protein labeling with small molecule fluorophores

To produce Cy3 labeled FUS LC, a FUS LC fusion protein with a Cys-Ser-Gly C-terminal tag (effectively FUS-LC(1-166) S164C) was generated by introducing the requisite sequence to the wild-type FUS LC plasmid using a NEBuilder^®^ HiFi DNA Assembly Cloning Kit. The 6xHis-MBP-FUS LC(1-166) S164C construct was expressed and subjected to Ni-NTA purification as described above. The eluent containing 6xHis-MBP-FUS LC(1-166) S164C was diluted to a final concentration of 20 μM with reaction buffer (20 mM sodium phosphate, 300 mM sodium chloride, pH 7.4), and 500 μM TCEP and 80 μM Cy3-maleamide (APExBIO, 4 molar eq.) were added. The reaction was allowed to proceed for 60 seconds at 25°C in the dark before quenching with excess β-mercaptoethanol (>200 molar eq). The reaction mixture was then transferred to 10 kDa MWCO dialysis tubing and dialyzed twice into 20 mM sodium phosphate, 300 mM sodium chloride, 100 μM TCEP, pH 7.4 at 4°C. Once the majority of the residual Cy3 was removed by dialysis, the labeled protein was removed from the dialysis tubing and subjected to TEV cleavage and size-exclusion chromatography as described above. The labeling efficiency was ~30% as determined by analytical high-pressure liquid chromatography (HPLC) and mass spectrometry.

### Microscopy of liquid-liquid phase separated droplets

Liquid-liquid phase separation was induced by the dilution of 1.8 mM stock solutions of FUS LC (with 5% FUS LC(1-166) S164C-Cy3) in 20 mM CAPS, 150 mM sodium chloride, pH 11 with 20 mM sodium phosphate, 150 mM sodium chloride, pH 7.4 at 25°C. Protein stocks were added to buffer to mitigate the risk of aggregation. After the addition of protein, phase separation was apparent by the sample rapidly becoming cloudy and opaque. The phase separated protein was added to a microscope slide and allowed to incubate quiescently over the course of the imaging experiment. In order to prevent both the evaporation of the buffer and the mechanical perturbation of the droplets, the borders of the microscope cover slips were coated with a small amount of vacuum grease that served to raise the slide and to hermetically seal the sample.

### Fluorescence recovery after photobleaching

Droplet samples were prepared as described above and imaged on an Olympus FV1000 Confocal microscope. Six droplets with 5-10 μM diameter were subjected to photobleaching within a circular region with a diameter representing half of the droplet. Photobleaching was performed for 2 seconds using an FV3000 Hybrid Scan Unit in Tornado Scanning mode. Fluorescence intensity was recorded within the bleached region every 2 seconds. Data were normalized to the pre- and post-bleach fluorescence. All microscopy images were processed with Fiji^49^/ImageJ^50^ and data were analyzed and visualized with SciPy tools^51–54^.

### Thioflavin T assays

Spectra were recorded at 25 °C on a Molecular Devices SpectraMax i3x fluorometer. The excitation wavelength was 440 nm and emission was recorded from 465 to 520 nm at a scan speed of 1 nm/s. Liquid-liquid phase separation was induced by the dilution of 1.8 mM stock solutions of FUS LC (stored in 20 mM CAPS, 150 mM sodium chloride, pH 11) with 20 mM sodium phosphate, 150 mM sodium chloride, pH 7.4 to a final concentration of 300 μM FUS LC. A phase separated stock solution was kept and an aliquot was drawn at each time point for analysis. Each aliquot contained 30 μM FUS LC and 20 μM thioflavin T diluted with 20 mM sodium phosphate, 150 mM sodium chloride, pH 7.4.

### Solid-state nuclear magnetic resonance experiments

All experiments were performed using 3.2 mm thin-walled zirconia MAS rotors with 50 μL sample volume. Liquid-liquid phase separation of 30 mg ^15^N, ^13^C-labeled FUS LC or ^15^N, ^13^C-labeled FUS LC G165E was induced as described above to 300 μM and the droplets were transferred into the rotor by gentle centrifugation at 3,000 ×*g* using a device built in-house. This condensed the sample into a single proteinaceous phase with final concentration of approximately 400 mg/ml (23 mM). This results in approximately 20 mg of protein inside the rotor. Spectra were acquired on a 750 MHz (17.6 T) NMR spectrometer equipped with a 3.2 mm E^free^ triple resonance HCN MAS probe (Bruker Biospin). All experiments were performed at MAS frequency of 11.11 kHz and 285 K. More details regarding the MAS NMR experimental settings are given in the Supplementary Information. Data were visualized and analyzed with NMRFAM-Sparky^55^.

### Statistical analysis of NMR chemical shifts

Chemical shift and protein coordinate data were acquired from the PDB and BMRB with PACSY Maker^56^. For each amino acid in FUS LC, the PACSY database was filtered using secondary structure classifications generated by STRIDE^57^ to produce a dataset of chemical shifts associated with residues in PDB structures that are in either β-sheet or random coil conformations. Only chemical shifts with unambiguous assignments were considered. Since the BMRB only contains 1D assignments, chemical shifts were projected into 2 dimensions by considering atom connectivity for each amino acid and plotting theoretical correlations using the chemical shifts for directly bonded atoms on a per-protein basis. Data were analyzed and results were visualized using SciPy tools^51–54^. Bruker NMR data were parsed with Nmrglue^58^. All code is available upon request. It should be noted that this approach is similar to that taken by PLUQ^59^.

### Coarse grained molecular dynamics simulations

Molecular dynamics simulations were performed with HOOMD-blue^60^ using a set of hydrophobicity-scaled pair potentials initially described by Dignon, *et al*^28^. This model has been shown to be effective for simulating liquid-liquid phase separation of FUS LC and other proteins^32,48,61,62^. Briefly, the model defines three interaction potentials–a potential for bonded interactions, a single potential representing short range non-bonded interactions, and a potential representing electrostatic interactions. Bonds are represented by a harmonic potential with a bond length of 3.8 Å and a spring constant of 10 kJ/Å^2^. Short range non-bonded interactions are represented by a standard Lennard-Jones potential that has been scaled to the hydrophobicity of each interaction pair. This scaled Lennard Jones potential was utilized as implemented in the azplugins package for HOOMD-blue^63^, and the hydrophobicity scaling parameters were identical to those described in Dignon, *et al*^28,64^. Electrostatic interactions were represented by the Yukawa potential with a Debye screening length of 1 nm and a dielectric constant of 80 to mimic an aqueous solvent containing 100 mM salt. For each simulation, 100 FUS LC or FUS LC G156E monomers were prepared in a linear configuration using mBuild^65^. The coordinates of each particle in each monomer were then randomized such that each bond length was fixed at 3.8 Å and no particles overlapped. Monomers were then packed into a 50 nm^3^ simulation cell using PACKMOL^66^. Charges and masses were assigned to each particle within HOOMD-blue. Production simulations were performed on a single NVIDIA Tesla K80 GPU. Each simulation was run for a total of 1.1 μs with 10 fs timesteps using a Langevin integrator at 300 K. Simulations were analyzed and contact maps were visualized using tools from the SciPy software stack^51–54^. Simulation snapshots were visualized using Ovito Pro^67^.

## Results and Discussion

### Liquid droplets of FUS LC undergo transition to gels and fibers at neutral pH

The full-length FUS protein can undergo a liquid-to-solid phase transition that is accelerated by the presence of a clinically relevant G156E mutation^19^. We chose to work with the N-terminal low complexity domain comprising residues 1-163 of the protein (FUS LC) to investigate the differential behavior of FUS LC and the pathological FUS LC G156E mutant *in vitro*. The low complexity domain of FUS has been extensively studied in the LLPS state^31,32,48^ and includes residues 39-95 that are known to form a rigid fibril core^16^. We prepared recombinantly purified protein (**Fig S1, Fig S2**) mixed with 5% Cy3-labeled FUS LC and initiated phase separation under physiological pH and low salt conditions. We observed the immediate formation of liquid-liquid phase separated droplets that were subsequently placed in hermetically sealed microscope slides and monitored for several weeks (**Fig 1A**). The well-defined droplets coarsened into gel-like structures over the course of several days. As the droplets continued to age, fibrillar structures developed that appeared to protrude from the gel cores in a manner consistent with previous reports for FUS and other proteins with low complexity sequences capable of undergoing liquid-to-solid phase transitions ^12,21^. After incubation for a full month, these fibrillar structures had grown into a dense network covering the microscope slide. Two-month old samples were also subjected to negative staining and transmission electron microscopy (TEM) where dense clumps of fibers were also observed (**Fig S3**). In contrast to wild-type FUS LC, the FUS LC G156E mutant formed fibrillar species much more rapidly, with clear fibrillar protrusions appearing after only a day of incubation (**Fig 1B**). Thus, the FUS LC and FUS LC G156E constructs recapitulate the transformation behavior of the full-length protein at neutral pH as described previously^19,68,69^.

**Figure 1:**
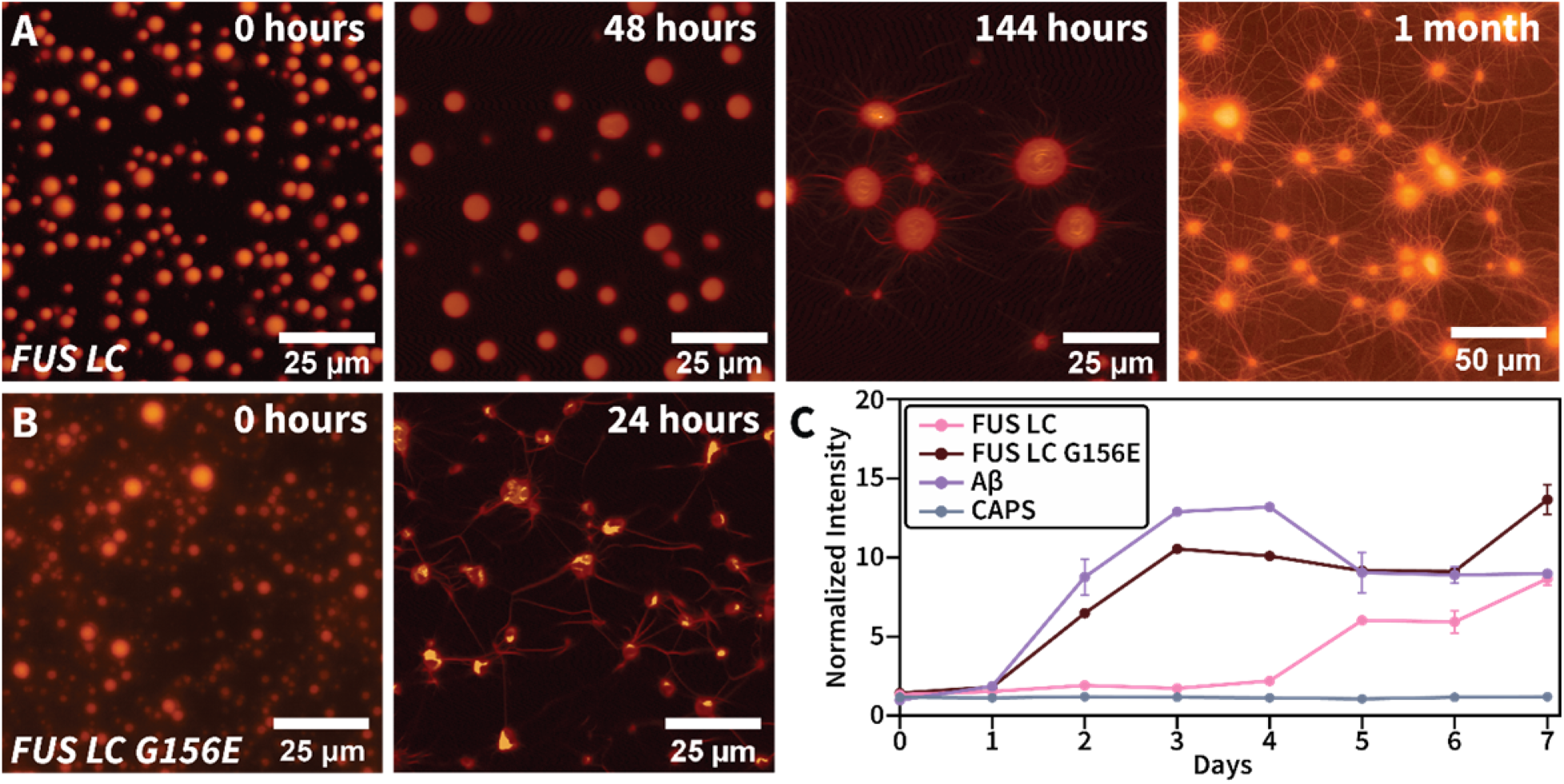
Microscopy and Thioflavin T (ThT) analysis of FUS LC and FUS LC G156E maturation. **A)** Maturation of wild-type FUS LC droplets. Droplets coarsen and loose spherical character after 48 hours. Fibrils begin to appear after 144 hours. After a full month, a dense fibril network is visible. **B)** Maturation of FUS LC G15E droplets. The progression of the G156E mutant is accelerated, with aggregates and fibrils appearing after 24 hours. C) Fibrillization rates of FUS LC and FUS LC G156E characterized by ThT. The A² peptide was used as positive control while FUS LC in CAPS buffer (pH 11) served as negative control. The error bars represent standard deviation from three independent measurements.

The maturation of FUS LC droplets was further interrogated by fluorescence recovery after photobleaching (FRAP) experiments (**Fig S4A**). Although the recovery kinetics of FUS LC droplets decreased over the course of several weeks, fluorescence recovery was not completely abrogated even after eight weeks, suggesting that mobile components remained in phase-separated droplets even after this long period of incubation. The FUS LC G156E sample also exhibited fast recovery at the beginning but transitioned quickly to a fibrillar state with limited mobility (**Fig S4B**). Interestingly, the initial recovery kinetics appear to be faster for the G156E mutant compared to the wild-type FUS LC sample (**Fig S4B**), suggesting faster reorganizational dynamics within that droplet environment.

The observation of fibrillar structures in our FUS LC and FUS LC G156E samples by microscopy prompted us to determine whether these structures have the characteristics of β-sheet rich amyloid fibers. Thioflavin T is a switchable small molecule fluorophore that gives a fluorescence signal upon binding to amyloid-like fibrils and is often used as a test for the presence of amyloid^70^. We initiated LLPS of FUS LC and FUS LC G156E samples and performed a binding assay with Thioflavin T (ThT) (**Fig 1C**). Consistent with the trends observed in the microscopy experiments, FUS LC G156E formed fibers within the first two days and with similar kinetics to a positive control sample containing the amyloidogenic Aβ peptide. On the other hand, ThT fluorescence of the wild-type FUS LC sample started to increase after approximately a week while a negative control sample containing solubilized FUS LC in high pH CAPS buffer did not form amyloid over the course of the experiment. These changes in fluorescence intensity are consistent with the timeline of visual changes observed in the microscopy samples. We also performed the experiment in the presence of 1,6-hexanediol, a small molecule known to disrupt protein droplets but not mature states such as gels and fibers (**Fig S5**)^71^. In this case, no ThT fluorescence was observed over the course of this experiment. Taken together, these data indicate that wild-type FUS LC droplets can coarsen into gel-like states and mature into amyloid fibrils over time under near-physiological conditions. For the FUS LC G156E mutant, the transition from droplets to gels is more abrupt and the formation of the amyloid state is significantly accelerated.

### The structure and dynamics of FUS LC change during the transformation process

The interactions and dynamics of FUS LC in the liquid droplet state have been characterized extensively by solution NMR^31,32,48^. On the other hand, solid-state MAS NMR spectroscopy has revealed the structure of the final amyloid state^16,35,36^. We sought to establish a connection between these two states by observing the transformation from droplet to amyloid on the molecular level in real time using MAS NMR. The maturation of FUS LC droplets into gels and fibrils is ideally suited to this approach as the transformation of the wild-type protein takes several weeks and thus allows sufficient time for the collection of multidimensional NMR experiments at different time points.

In order to capture the range of dynamic regimes within the sample, we performed two different types of MAS NMR experiments. First, the INEPT-based pulse sequence was used to capture mobile components in the sample. This can include the mobile segments of an otherwise slow tumbling protein system or the mobile subpopulation of proteins in a heterogeneous sample^16,47,72,73^. Second, dipolar-based experiments such as cross-polarization (CP) were used to describe the slow tumbling (rigid) components of the sample like those subpopulations in the gel and amyloid state^74,75^. As control, we also used a direct polarization (DP) ^13^C experiment which detects all carbon atoms in the sample irrespective of their mobility. We prepared two samples, one of ^15^N,^13^C FUS LC and one of ^15^N,^13^C FUS LC G156E, and subjected each to LLPS. The LLPS droplets were then gently collapsed into a single condensed phase and into a MAS NMR rotor. The final concentration of both FUS LC constructs was approximately 23 mM; within the range of the expected concentration of FUS LC in LLPS droplets^32^. We used INEPT, CP, and DP experiments to follow the transformation of the samples over the course of 30 days for FUS LC and 12 days for FUS LC G156E (**Fig 2**).

**Figure 2:**
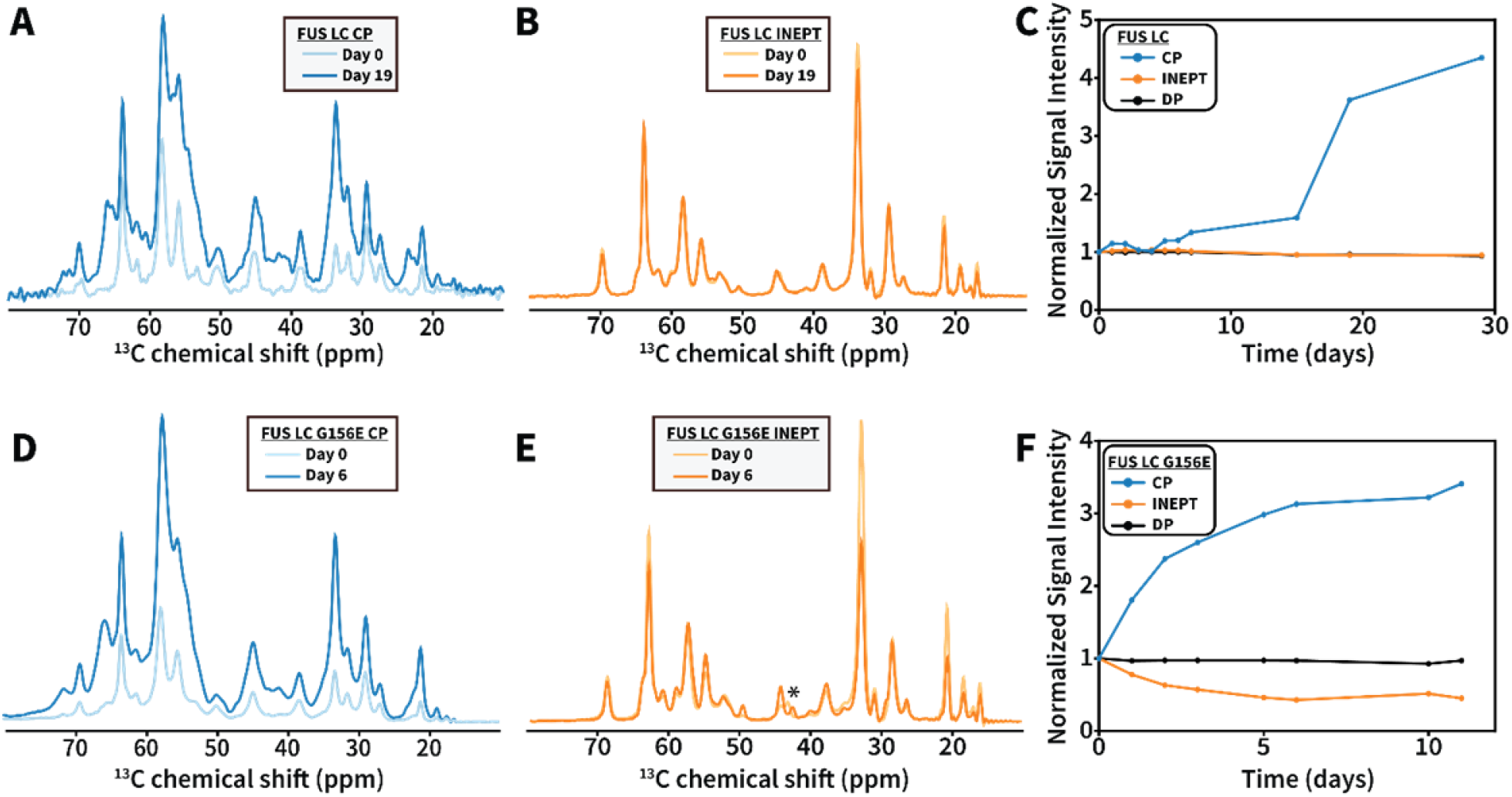
Characterization of FUS LC and FUS LC G156E maturation by MAS NMR spectroscopy. **A)** Early and late-stage FUS LC CP spectra. **B)** Early and late-stage FUS LC INEPT spectra. **C)** Integrated signal intensity of the aliphatic region of FUS LC over time for CP, INEPT and DP experiments. The error bars are based on the integrated noise level and are too small to visualize. **D)** Early and late-stage FUS LC G156E CP spectra. **E)** Early and late-stage FUS LC G156E INEPT spectra. The asterisks denotes a region of the spectrum where changes are observed over time. **F)** Integrated signal intensity of the aliphatic region of FUS LC G156E over time for CP, INEPT and DP experiments. The error bars are based on the integrated noise level and are too small to visualize.

The two samples exhibited different behavior over the course of these experiments. For the wild-type FUS LC sample, the CP signal started to increase after a week, eventually reaching a plateau at 3.5 times the original integrated signal (**Fig 2A, C**). This indicates the emergence of rigid components in the sample with a timeline that is consistent with the formation of the fibrillar species detected by microscopy and ThT fluorescence. Furthermore, a comparison of the initial and final CP spectra shows the appearance of several new peaks (**Fig 2A, S6A, B**). This includes peaks consistent with the chemical shifts of threonine, serine, glycine and tyrosine. These peaks represent new chemical environments and suggest that the rigid components in the sample undergo a structural change over time, perhaps from an unstructured or oligomeric state to a β-sheet rich amyloid fold. In contrast, the FUS LC G156E sample changed much more rapidly over the course of a week (**Fig 2D, F**). A comparison of the CP spectra shows a 3-fold increase in signal and the appearance of some new structural environments during this time (**Fig S6C, D**). Overall, however, the changes in structure are not as pronounced as those for the wild-type sample suggesting that a significant population of the proteins in this sample has already adopted a rigid core structure at the early stages of the experiment.

The time course of the INEPT experiments is also noteworthy. While the INEPT spectra of the wild-type protein changed very little over the course of three weeks (**Fig 2B, C**), the signals of the G156E sample decreased significantly during the first week, and then stabilized at ~50% of the initial intensity (**Fig 2E, F**). In these heterogenous samples, the INEPT signals potentially arise from two different processes. First, they may reflect mobile monomers or low molecular weight oligomers that experience fast diffusion and rotational correlation times within the phase separated liquid or gel environment that surrounds the nascent amyloid fibers. These mobile components should also be detectable in FRAP experiments and, as our data indicate, their contribution diminishes over time (**Fig S4**). Second, these signals may arise from mobile regions of the protein sequence. For example, the published structure of seeded amyloid fibers of FUS LC(1-214) has a rigid β-sheet core that spans 57 residues (39-95), while the rest of the sequence remains dynamic and unstructured^16^. This type of dynamic behavior would be invisible in the FRAP experiments and is expected to increase during the transformation process. It should also be noted that INEPT often does not capture the full range of motions in the sample as sometimes these motions can interfere with decoupling during acquisition or might be in an intermediate regime that causes line broadening. While much more detailed and targeted work is needed to untangle the relative contributions of these processes, it is clear that the G156E sample experiences much more profound changes in mobility over the course of the experiment as detected by INEPT. More interestingly, however, there are also small but detectable chemical shift differences between the initial and final INEPT spectra of this sample (**Fig 2E**). These changes occur around 45 ppm which is the glycine Cα region of the spectrum, an observation that will be discussed in more detail below.

### LLPS results in amyloid fibers with distinct structures

Intrigued by the structural changes detected in the 1D MAS NMR experiments described above, we recorded 2D correlation spectra of the end-states of the two samples (30 days for FUS LC and 12 days for FUS LC G156E). In particular, we extended the 1D INEPT experiment into a 2D ^1^H-^13^C INEPT correlation spectrum that provides a more detailed picture of the residues that remain mobile at the end of the time course (**Fig 3A**). The 2D INEPT spectra of the wild-type and G156E samples contain similar amino acids types including glycine, threonine, serine, glutamine, alanine, proline and methionine with chemical shifts that are consistent with random coil. The INEPT spectrum of the G156E construct has lower intensity overall despite the comparable number of scans and sample amount in the rotor, consistent with the 50% reduction of INEPT signal observed over the course of the 1D experiments. Despite the general overlap between the two spectra, a careful comparison reveals differences in the glycine region where one set of glycine residues remains the same while another set experiences a different chemical environment. This matches the new glycine peak observed in the 1D spectra above (**Fig 2E)**. Due to the large number of glycine residues in the FUS sequence, we were not able to assign these two different glycine groups. Since they are present at the very end of the time course when the sample is no longer changing, it is possible that they represent two different mobile glycine environments within the FUS LC G156E fibrils. However, we also cannot exclude the possibility that they arise from two different components in the heterogeneous sample, e.g. dynamic segments in the fibril structure and mobile monomers within the gel environment encapsulating those fibers. Overall, however, there appear to be subtle differences between the dynamic environments in the wild-type and mutant FUS LC samples.

**Figure 3:**
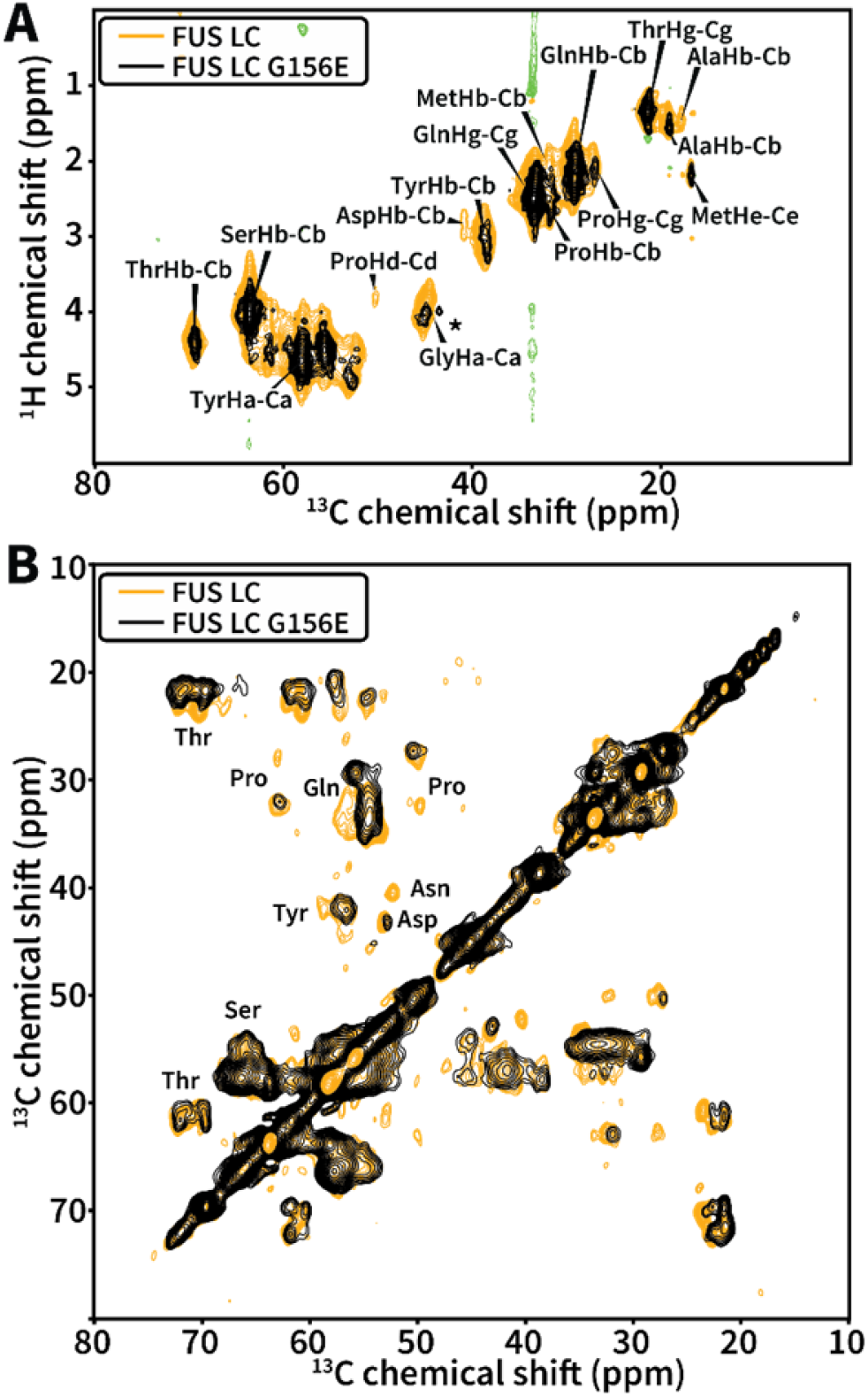
2D correlation spectra of FUS LC and FUS LC G156E. **A)** End-state ^1^H-^13^C INEPT spectra of FUS LC (orange) and FUS LC G156E (black). The asterisks denotes a new glycine cross-peak that appears in the FUS LC G156E sample. **B)** End-state ^13^C-^13^C DARR correlation spectra of FUS LC (orange) and FUS LC G156E (black).

To complement the residue specific studies of mobile sample components, we also recorded 2D 13C-^13^C DARR spectra of the final state (**Fig 3B**). This experiment relies on a dipolar assisted rotational resonance condition to recouple ^13^C atoms in protein regions that experience reduced motions and rotational correlation times^76^. Given the composition of the two samples, this experiment is expected to report on the rigid core of the final amyloid state. Both samples gave rise to relatively well resolved DARR spectra where many individual cross peaks could be identified and analyzed. In order to determine whether the spectra indeed report on a β-sheet rich amyloid fold, we compared the position of the cross-peaks against a statistical analysis of chemical shifts deposited in the Biological Magnetic Resonance Bank (BMRB) and the Protein Data Bank (PDB) (**Fig S7**). Plotting these distributions with our DARR spectrum reveals that many of the correlations in our DARR lie nearer to the center of the β-sheet distribution than the coil-like distribution (**Fig S7**). These observations suggest that the rigid structures appearing in our DARR spectrum are β-sheet-like.

Unlike previous studies of FUS LC fibrils^16^, which used seeded fibers in isolation, our samples are a heterogeneous mixture of gel-like and fibrillar states. Therefore, the signal-to-noise in the dipolar experiments is relatively low and this precluded the collection of other multidimensional MAS NMR correlations (e.g. 3D NCACX and 3D NCOCX). While this prevented us from performing site-specific assignments of the DARR spectra, we were able to compare our data with the DARR spectra of the published FUS LC(1-214) structure of seeded amyloid fibers (**Fig S8**). This comparison revealed significant differences between the two spectra. First, our spectrum is relatively broader, indicating the presence of heterogeneity and/or intermediate dynamics that can interfere with the timescale of the NMR experiment and cause line broadening. This is expected as our sample is more heterogenous and dynamic by design. Second, a significant number of cross peaks appear to be shifted or missing from our spectrum. This includes cross peaks for the unique residues Asp46 and Pro72 that appear to be substantially shifted, as well as cross peaks for residues such as Gln93 that are not present altogether. Other residues that potentially experience different environments or are not part of the core include Thr45, Thr47, Ser70, Thr78 and Ser90. Based on this analysis, it appears that the core of LLPS-derived fibrils is in a similar region of the FUS LC sequence and includes Asp46 and Pro72. However, the structure of the core is different as manifested by the significant chemical shift changes of the cross-peaks. It should also be noted that our DARR spectrum contains a much more intense tyrosine Cα-Cβ region potentially implying a more prominent role for these residues in the structure and dynamics of the final state.

Finally, we compared the DARR spectra of the wild-type FUS LC and the G156E mutant (**Fig 3B**). The two spectra are much more similar to each other than to the DARR spectrum of the seeded FUS LC(1-214) amyloids. This includes key residues such as Pro72 and Asp46 which have similar conformations in our samples, while other residues such as Gln93 are prominently missing from our spectra. The combined analyses of the INEPT and DARR spectra indicate that LLPS-derived amyloid fibrils have a distinct fold compared to fibers obtained through seeding, and that the structure and dynamics of the fibers are further fine-tuned by clinically relevant mutations such as the G156E mutation investigated here.

### The G156E mutation increases the contact propensity of FUS LC

Previous work has shown that phosphorylation of serine and threonine residues in the FUS LC domain can inhibit LLPS^16,48^. Incubation at high pH also disrupts LLPS likely due to the deprotonation and subsequent negative charge of tyrosine side chains in FUS LC under these conditions^31^. It is therefore noteworthy that the G156E mutant, which introduces a phosphomimetic, negatively charged residue into FUS LC, can not only undergo LLPS but also exhibits faster aggregation kinetics than the wild type. Furthermore, based on the published structure of seeded fibrils^16^ and our MAS NMR analysis, this residue does not appear to participate in the ordered fibril core. Therefore, it is still unclear how this mutation promotes amyloidogenesis from an LLPS state. To gain insight into the early events that may drive this behavior, we used coarse-grained molecular dynamics simulations to model interactions in liquid droplets formed by wild-type FUS LC and the FUS LC G156E mutant. We applied a hydrophobicity scale model^64^, initially described by Dignon et al., which is capable of recapitulating LLPS *in silico* and has been used to analyze interactions in wild-type FUS, FUS LC, and other phosphomimic mutants of FUS LC^28,32^.

Separate simulations of 100 dispersed FUS LC or FUS LC G156E monomers in random configurations were prepared and allowed to proceed for 1 μs. In both simulations, dispersed monomers rapidly coalesced into a dynamic droplet-like state within 30 ns. The droplet persisted throughout the simulation (**Fig S9**), with some monomers occasionally breaking free from the LLPS droplet and rejoining after a few ns. We analyzed the coalesced state by constructing intermolecular contact maps that represent the number of pair-wise amino acid contacts within a specified radius. For each simulation run, 100 frames from the droplet state were extracted and interparticle distances were measured for all particles in the simulation. Particles that were within the specified distance radius were counted as a contact. Intramolecular (i.e. intramonomer) interactions were not counted. The mean number of contacts across all frames and monomers was used to construct the contact maps.

In alignment with previous simulations of FUS LC^32^, there are no notable regions of high contact propensity within a contact radius of up to 7 Å (**Fig S10**). This is also consistent with experimental literature that suggests that LLPS of FUS LC is not driven by specific regions in the sequence^32^. At longer contact distances (8 – 10 Å), however, some features started to appear in the contact map (**Fig 4, S10**). This includes fewer contacts between the regions surrounding the two native negatively charged residues, Asp5 and Asp46, as well as increased contacts between a segment encompassing residues 15-25 and another segment comprising the more aliphatic region of FUS LC between residues 99 – 110. It is noteworthy that those interacting regions flank the rigid fibril core of the seeded fibers (39-95) but are not part of it. The increased propensity for contacts in this region was also evident in the FUS LC G156E simulation. These features are interesting given that interactions on this length scale are relevant for amyloidogenesis, as the characteristic β-sheet amyloid structures have backbone to backbone distances of ~10 Å.

**Figure 4:**
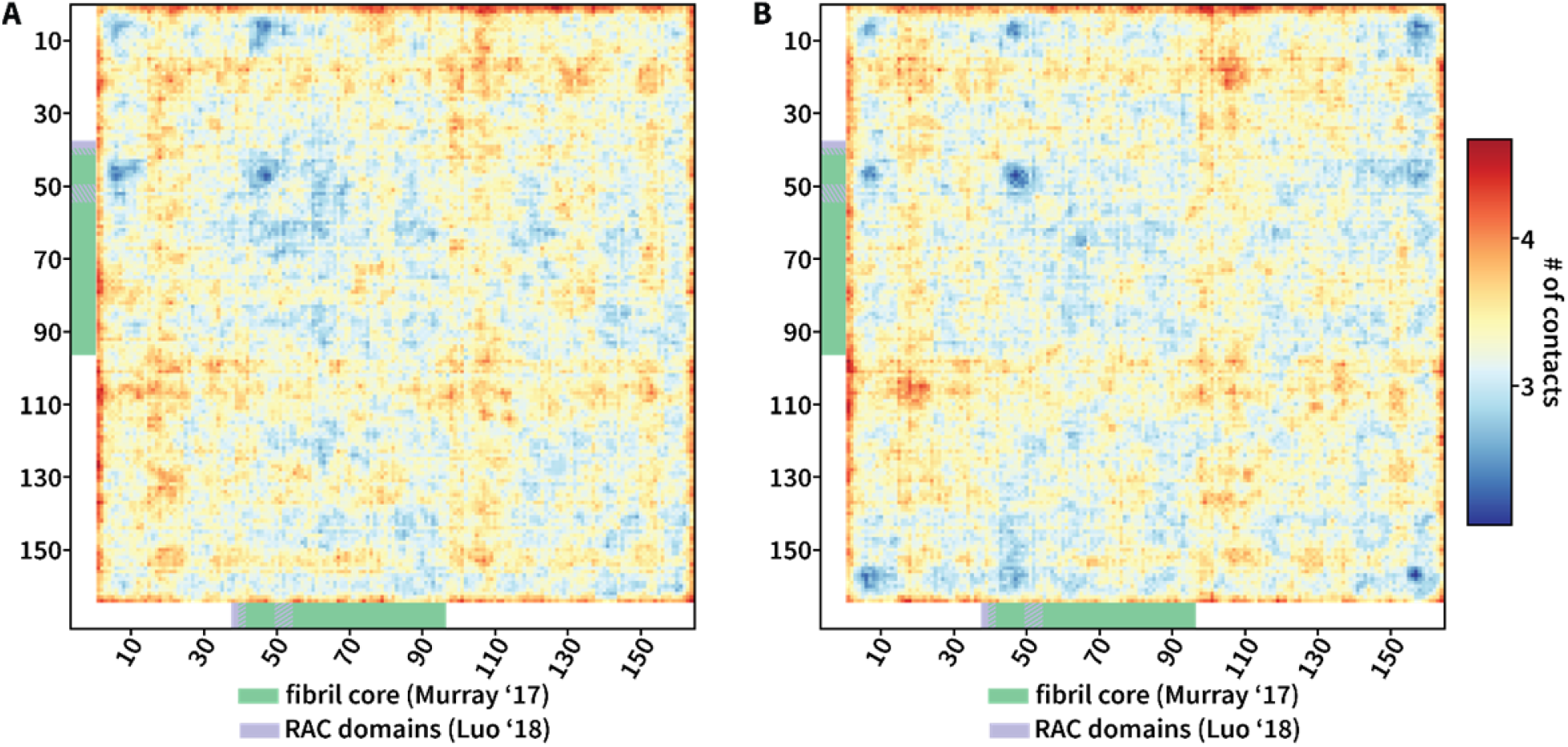
Coarse-grained simulations of FUS LC and FUS LC G156E droplets. **A)** Intermolecular contact map for FUS LC. Regions with a large number of contacts appear in orange, while regions with relatively low number of contacts appear in blue. **B)** Intermolecular contact map for FUS LC G156E. The color coding for both maps is the same and both maps represent contacts within 10 Å cutoff. Distance is measured from the bead centers.

The primary difference between the FUS LC and FUS LC G156E contact maps lies near the terminal regions (**Fig 4B**). The introduction of a negative charge at position 156 decreases the number of contacts between the C-terminus and the N-terminal Asp5. Fewer contacts are also observed between Glu156 and Asp46. Overall, the repulsion between these negatively charged residues reduces the interactions of the C-terminus with the majority of the N-terminal part of the protein. At the same time, the higher contact propensities of regions 15-25 and 99-110 are preserved. In light of these observations, we propose that the introduction of the G156E mutation into FUS LC reduces the number of non-productive interactions between the termini and leads to greater exposure of the inner segments of the protein. This increases the propensity for protein-protein interactions that, in turn, leads to amyloid formation on an accelerated timescale.

## Conclusions

The low complexity domain of FUS can undergo phase transitions that lead to the formation of liquid droplets, gels, and amyloid. Our work demonstrates that that FUS LC droplets formed at neutral pH mature into amyloid-like states over the course of several weeks, a transition that can be significantly accelerated by the disease-relevant mutation G156E. The slow kinetics of the process allowed us to follow this transition by MAS NMR spectroscopy and to observe the formation of characteristic β-sheet signatures at residue specific resolution in real time. While the resulting β-sheet core of these LLPS-derived fibrils is located in the same region of the sequence as the rigid core formed by seeded fibers, the core structure of the LLPS-derived fibrils is substantially different. Furthermore, when the LLPS environment was eliminated by the addition of 1,6-hexanediol, no amyloid fibrils were observed in the course of the experiment. These observations imply that liquid droplet environments can influence both the efficiency of fibril formation and their structural signatures. Considering that FUS is often located in liquid droplet like stress granules in the cell^77^, the crowded conditions promoted by LLPS may play an important role in its pathogenic amyloid formation.

In contrast to the differences observed between the fibril core structure of seeded FUS LC fibrils and LLPS-derived fibrils, the fibril core structures of the LLPS-derived FUS LC and FUS LC G156E constructs appear relatively similar to each other. The most striking differences between the wild-type protein and the pathological mutant were maturation rate rather than the final structure, with the FUS LC G156E protein forming fibrils much earlier than its wild-type counterpart under the same conditions. While the wild-type FUS LC sample has a significant mobile component even after several weeks, the G156E mutant matures much more rapidly and loses mobility over the course of several days. Thus, a single phosphomimic mutation at a region of the sequence that does not participate in fibril core interactions can have dramatic consequences on the fibrillization process and the dynamics of the resulting fiber. Our coarse-grained simulations suggest that the presence of this negatively charged residue near the C-terminus increases the repulsion between the termini and promotes enhanced intermolecular contacts between inner regions of the protein. This mechanism may also explain the increased aggregation propensity of the full-length G156E mutant^19^.

Our approach presented here combines imaging, MAS NMR spectroscopy and computational methods to provide new molecular insights into the elusive transformation of a protein from the liquid droplet to the gel and amyloid states. Our strategy is versatile and provides a platform for the analysis of other proteins that exhibit similar behavior in viscous and heterogeneous environments. It can also be extended to explore the role of relevant biological components and small molecules on phase separation, gelation and amyloid formation.

## Author Contributions

R.F.B. and G.T.D. conceived the project and wrote the manuscript. R.F.B. prepared samples, performed microscopy, ThT assays, NMR experiments, and computational studies. M.K. performed ThT assays and NMR experiments. G.T.D. performed NMR experiments and supervised the study. All authors analyzed the data and commented on the manuscript.

## Acknowledgments

We are grateful to A. de Angelis for help with the NMR spectrometers, to J. Yang’s lab at UCSD for help with the ThT assays, to T. Meerloo for help with TEM, and to J. Mittal for generously sharing simulations code. This work was supported by a Research Education Component associated with NIH grant P30 AG062429, an R21 AG069064 award to G.T.D., a T32 GM112584 fellowship to R.F.B., the BRC for NMR Molecular Imaging of Proteins at UCSD (P41 EB002031) and the UCSD Microscopy Core (NINDS NS047101).

## References

1. Soto, C. & Pritzkow, S. Protein misfolding, aggregation, and conformational strains in neurodegenerative diseases. Nat. Neurosci. 21, 1332–1340 (2018).

2. Mathieu, C., Pappu, R. V. & Taylor, J. P. Beyond aggregation: Pathological phase transitions in neurodegenerative disease. Science 370, 56–60 (2020).

3. Wang, J. et al. A Molecular Grammar Governing the Driving Forces for Phase Separation of Prion-like RNA Binding Proteins. Cell 174, 688–699.e16 (2018).

4. Martin, E. W. et al. Valence and patterning of aromatic residues determine the phase behavior of prion-like domains. Science 367, 694–699 (2020).

5. Fitzpatrick, A. W. P. et al. Cryo-EM structures of tau filaments from Alzheimer’s disease. Nature 547, 185–190 (2017).

6. Lin, Y., Fichou, Y., Zeng, Z., Hu, N. Y. & Han, S. Electrostatically Driven Complex Coacervation and Amyloid Aggregation of Tau Are Independent Processes with Overlapping Conditions. ACS Chem. Neurosci. 11, 615–627 (2020).

7. Wegmann, S. et al. Tau protein liquid–liquid phase separation can initiate tau aggregation. EMBO J. 37, e98049 (2018).

8. Dregni, A. J. et al. In vitro 0N4R tau fibrils contain a monomorphic β-sheet core enclosed by dynamically heterogeneous fuzzy coat segments. Proc. Natl. Acad. Sci. 116, 16357–16366 (2019).

9. Ambadipudi, S., Reddy, J. G., Biernat, J., Mandelkow, E. & Zweckstetter, M. Residue-specific identification of phase separation hot spots of Alzheimer’s-related protein tau. Chem. Sci. 10, 6503–6507 (2019).

10. Ambadipudi, S., Biernat, J., Riedel, D., Mandelkow, E. & Zweckstetter, M. Liquid–liquid phase separation of the microtubule-binding repeats of the Alzheimer-related protein Tau. Nat. Commun. 8, 1–13 (2017).

11. Schweighauser, M. et al. Structures of α-synuclein filaments from multiple system atrophy. Nature 1–6 (2020) doi:10.1038/s41586-020-2317-6.

12. Ray, S. et al. α-Synuclein aggregation nucleates through liquid–liquid phase separation. Nat. Chem. 12, 705–716 (2020).

13. Araki, K. et al. Parkinson’s disease is a type of amyloidosis featuring accumulation of amyloid fibrils of α-synuclein. Proc. Natl. Acad. Sci. 201906124 (2019) doi:10.1073/pnas.1906124116.

14. Tuttle, M. D. et al. Solid-state NMR structure of a pathogenic fibril of full-length human α-synuclein. Nat. Struct. Mol. Biol. 23, 409–415 (2016).

15. Li, Y. R., King, O. D., Shorter, J. & Gitler, A. D. Stress granules as crucibles of ALS pathogenesis. J. Cell Biol. 201, 361–372 (2013).

16. Murray, D. T. et al. Structure of FUS Protein Fibrils and Its Relevance to Self-Assembly and Phase Separation of Low-Complexity Domains. Cell 171, 615–627.e16 (2017).

17. Gui, X. et al. Structural basis for reversible amyloids of hnRNPA1 elucidates their role in stress granule assembly. Nat. Commun. 10, 1–12 (2019).

18. Babinchak, W. M. et al. The role of liquid–liquid phase separation in aggregation of the TDP-43 low-complexity domain. J. Biol. Chem. 294, 6306–6317 (2019).

19. Patel, A. et al. A Liquid-to-Solid Phase Transition of the ALS Protein FUS Accelerated by Disease Mutation. Cell 162, 1066–1077 (2015).

20. Lin, Y., Protter, D. S. W., Rosen, M. K. & Parker, R. Formation and Maturation of Phase-Separated Liquid Droplets by RNA-Binding Proteins. Mol. Cell 60, 208–219 (2015).

21. Liu, Z. et al. Hsp27 chaperones FUS phase separation under the modulation of stress-induced phosphorylation. Nat. Struct. Mol. Biol. 27, 363–372 (2020).

22. Mitrea, D. M. et al. Methods for Physical Characterization of Phase-Separated Bodies and Membrane-less Organelles. J. Mol. Biol. 430, 4773–4805 (2018).

23. Mittag, T. & Forman-Kay, J. D. Atomic-level characterization of disordered protein ensembles. Curr. Opin. Struct. Biol. 17, 3–14 (2007).

24. Martin, E. W. & Mittag, T. Relationship of Sequence and Phase Separation in Protein Low-Complexity Regions. Biochemistry 57, 2478–2487 (2018).

25. van der Lee, R. et al. Classification of Intrinsically Disordered Regions and Proteins. Chem. Rev. 114, 6589–6631 (2014).

26. Brangwynne, C. P., Tompa, P. & Pappu, R. V. Polymer physics of intracellular phase transitions. Nat. Phys. 11, 899–904 (2015).

27. Dignon, G. L., Best, R. B. & Mittal, J. Biomolecular Phase Separation: From Molecular Driving Forces to Macroscopic Properties. Annu. Rev. Phys. Chem. 71, 53–75 (2020).

28. Dignon, G. L., Zheng, W., Kim, Y. C., Best, R. B. & Mittal, J. Sequence determinants of protein phase behavior from a coarse-grained model. PLOS Comput. Biol. 14, e1005941 (2018).

29. Pak, C. W. et al. Sequence Determinants of Intracellular Phase Separation by Complex Coacervation of a Disordered Protein. Mol. Cell 63, 72–85 (2016).

30. Feric, M. et al. Coexisting Liquid Phases Underlie Nucleolar Subcompartments. Cell 165, 1686–1697 (2016).

31. Burke, K. A., Janke, A. M., Rhine, C. L. & Fawzi, N. L. Residue-by-Residue View of In Vitro FUS Granules that Bind the C-Terminal Domain of RNA Polymerase II. Mol. Cell 60, 231–241 (2015).

32. Murthy, A. C. et al. Molecular interactions underlying liquid−liquid phase separation of the FUS low-complexity domain. Nat. Struct. Mol. Biol. 26, 637 (2019).

33. Molliex, A. et al. Phase Separation by Low Complexity Domains Promotes Stress Granule Assembly and Drives Pathological Fibrillization. Cell 163, 123–133 (2015).

34. Banani, S. F. et al. Compositional Control of Phase-Separated Cellular Bodies. Cell 166, 651–663 (2016).

35. Murray, D. T. & Tycko, R. Side Chain Hydrogen-Bonding Interactions within Amyloid-like Fibrils Formed by the Low-Complexity Domain of FUS: Evidence from Solid State Nuclear Magnetic Resonance Spectroscopy. Biochemistry 59, 364–378 (2020).

36. Ding, X. et al. Amyloid-Forming Segment Induces Aggregation of FUS-LC Domain from Phase Separation Modulated by Site-Specific Phosphorylation. J. Mol. Biol. 432, 467–483 (2020).

37. Iadanza, M. G. et al. The structure of a β 2-microglobulin fibril suggests a molecular basis for its amyloid polymorphism. Nat. Commun. 9, 4517 (2018).

38. Ader, C. et al. Amyloid-like interactions within nucleoporin FG hydrogels. Proc. Natl. Acad. Sci. 107, 6281–6285 (2010).

39. Kennedy, S. B. et al. Dynamic Structure of a Protein Hydrogel: A Solid-State NMR Study. Macromolecules 34, 8675–8685 (2001).

40. Mandala, V. S. & Hong, M. High-sensitivity protein solid-state NMR spectroscopy. Curr. Opin. Struct. Biol. 58, 183–190 (2019).

41. Marchanka, A., Simon, B., Althoff-Ospelt, G. & Carlomagno, T. RNA structure determination by solid-state NMR spectroscopy. Nat. Commun. 6, 7024 (2015).

42. Andrew, E. R., Bradbury, A. & Eades, R. G. Nuclear Magnetic Resonance Spectra from a Crystal rotated at High Speed. Nature 182, 1659–1659 (1958).

43. Maricq, M. M. & Waugh, J. S. NMR in rotating solids. J. Chem. Phys. 70, 3300–3316 (1979).

44. Baldus, M. & Meier, B. H. Total Correlation Spectroscopy in the Solid State. The Use of Scalar Couplings to Determine the Through-Bond Connectivity. J. Magn. Reson. A 121, 65–69 (1996).

45. Takegoshi, K., Nakamura, S. & Terao, T. 13C–1H dipolar-assisted rotational resonance in magic-angle spinning NMR. Chem. Phys. Lett. 344, 631–637 (2001).

46. Matlahov, I. & van der Wel, P. C. A. Hidden motions and motion-induced invisibility: Dynamics-based spectral editing in solid-state NMR. Methods 148, 123–135 (2018).

47. Ackermann, B. E. & Debelouchina, G. T. Heterochromatin Protein HP1α Gelation Dynamics Revealed by Solid-State NMR Spectroscopy. Angew. Chem. Int. Ed. 58, 6300–6305 (2019).

48. Monahan, Z. et al. Phosphorylation of the FUS low-complexity domain disrupts phase separation, aggregation, and toxicity. EMBO J. 36, 2951–2967 (2017).

49. Schindelin, J. et al. Fiji: an open-source platform for biological-image analysis. Nat. Methods 9, 676–682 (2012).

50. Rueden, C. T. et al. ImageJ2: ImageJ for the next generation of scientific image data. BMC Bioinformatics 18, 529 (2017).

51. Virtanen, P. et al. SciPy 1.0: fundamental algorithms for scientific computing in Python. Nat. Methods 17, 261–272 (2020).

52. Harris, C. R. et al. Array programming with NumPy. Nature 585, 357–362 (2020).

53. McKinney, W. Data Structures for Statistical Computing in Python. Proc. 9th Python Sci. Conf. 56–61 (2010) doi:10.25080/Majora-92bf1922-00a.

54. Hunter, J. D. Matplotlib: A 2D Graphics Environment. Comput. Sci. Eng. 9, 90–95 (2007).

55. Lee, W., Westler, W. M., Bahrami, A., Eghbalnia, H. R. & Markley, J. L. PINE-SPARKY: graphical interface for evaluating automated probabilistic peak assignments in protein NMR spectroscopy. Bioinformatics 25, 2085–2087 (2009).

56. Lee, W. et al. PACSY, a relational database management system for protein structure and chemical shift analysis. J. Biomol. NMR 54, 169–179 (2012).

57. Heinig, M. & Frishman, D. STRIDE: a web server for secondary structure assignment from known atomic coordinates of proteins. Nucleic Acids Res. 32, W500–W502 (2004).

58. Helmus, J. J. & Jaroniec, C. P. Nmrglue: an open source Python package for the analysis of multidimensional NMR data. J. Biomol. NMR 55, 355–367 (2013).

59. Fritzsching, K. J., Yang, Y., Schmidt-Rohr, K. & Hong, M. Practical use of chemical shift databases for protein solid-state NMR: 2D chemical shift maps and amino-acid assignment with secondary-structure information. J. Biomol. NMR 56, 155–167 (2013).

60. Anderson, J. A., Glaser, J. & Glotzer, S. C. HOOMD-blue: A Python package for high-performance molecular dynamics and hard particle Monte Carlo simulations. Comput. Mater. Sci. 173, 109363 (2020).

61. Schuster, B. S. et al. Identifying sequence perturbations to an intrinsically disordered protein that determine its phase-separation behavior. Proc. Natl. Acad. Sci. 117, 11421–11431 (2020).

62. Conicella, A. E. et al. TDP-43 α-helical structure tunes liquid–liquid phase separation and function. Proc. Natl. Acad. Sci. 117, 5883–5894 (2020).

63. mphowardlab/azplugins. (mphowardlab, 2020).

64. Kapcha, L. H. & Rossky, P. J. A Simple Atomic-Level Hydrophobicity Scale Reveals Protein Interfacial Structure. J. Mol. Biol. 426, 484–498 (2014).

65. Klein, C. et al. A Hierarchical, Component Based Approach to Screening Properties of Soft Matter. in Foundations of Molecular Modeling and Simulation: Select Papers from FOMMS 2015 (eds. Snurr, R. Q., Adjiman, C. S. & Kofke, D. A.) 79–92 (Springer, 2016). doi:10.1007/978-981-10-1128-3_5.

66. Martínez, L., Andrade, R., Birgin, E. G. & Martínez, J. M. PACKMOL: A package for building initial configurations for molecular dynamics simulations. J. Comput. Chem. 30, 2157–2164 (2009).

67. Stukowski, A. Visualization and analysis of atomistic simulation data with OVITO–the Open Visualization Tool. Model. Simul. Mater. Sci. Eng. 18, 015012 (2009).

68. Murakami, T. et al. ALS/FTD Mutation-Induced Phase Transition of FUS Liquid Droplets and Reversible Hydrogels into Irreversible Hydrogels Impairs RNP Granule Function. Neuron 88, 678–690 (2015).

69. Nomura, T. et al. Intranuclear Aggregation of Mutant FUS/TLS as a Molecular Pathomechanism of Amyotrophic Lateral Sclerosis. J. Biol. Chem. 289, 1192–1202 (2014).

70. Naiki, H., Higuchi, K., Hosokawa, M. & Takeda, T. Fluorometric determination of amyloid fibrils in vitro using the fluorescent dye, thioflavine T. Anal. Biochem. 177, 244–249 (1989).

71. Kroschwald, S., Maharana, S. & Simon, A. Hexanediol: a chemical probe to investigate the material properties of membrane-less compartments. Matters 3, e201702000010 (2017).

72. Andronesi, O. C. et al. Determination of Membrane Protein Structure and Dynamics by Magic-Angle-Spinning Solid-State NMR Spectroscopy. J. Am. Chem. Soc. 127, 12965–12974 (2005).

73. Debelouchina, G. T., Platt, G. W., Bayro, M. J., Radford, S. E. & Griffin, R. G. Magic Angle Spinning NMR Analysis of β2-Microglobulin Amyloid Fibrils in Two Distinct Morphologies. J. Am. Chem. Soc. 132, 10414–10423 (2010).

74. Hartmann, S. R. & Hahn, E. L. Nuclear Double Resonance in the Rotating Frame. Phys. Rev. 128, 2042–2053 (1962).

75. Pines, A., Gibby, M. G. & Waugh, J. S. Proton-Enhanced Nuclear Induction Spectroscopy. A Method for High Resolution NMR of Dilute Spins in Solids. J. Chem. Phys. 56, 1776–1777 (1972).

76. Takegoshi, K., Nakamura, S. & Terao, T. 13C–1H dipolar-driven 13C–13C recoupling without 13C rf irradiation in nuclear magnetic resonance of rotating solids. J. Chem. Phys. 118, 2325–2341 (2003).

77. Wheeler, J. R., Matheny, T., Jain, S., Abrisch, R. & Parker, R. Distinct stages in stress granule assembly and disassembly. eLife 5, e18413 (2016).

