## Supplemental information for "Real-Time Observation of Structure and Dynamics during the Liquid-to-Solid Transition of FUS LC"

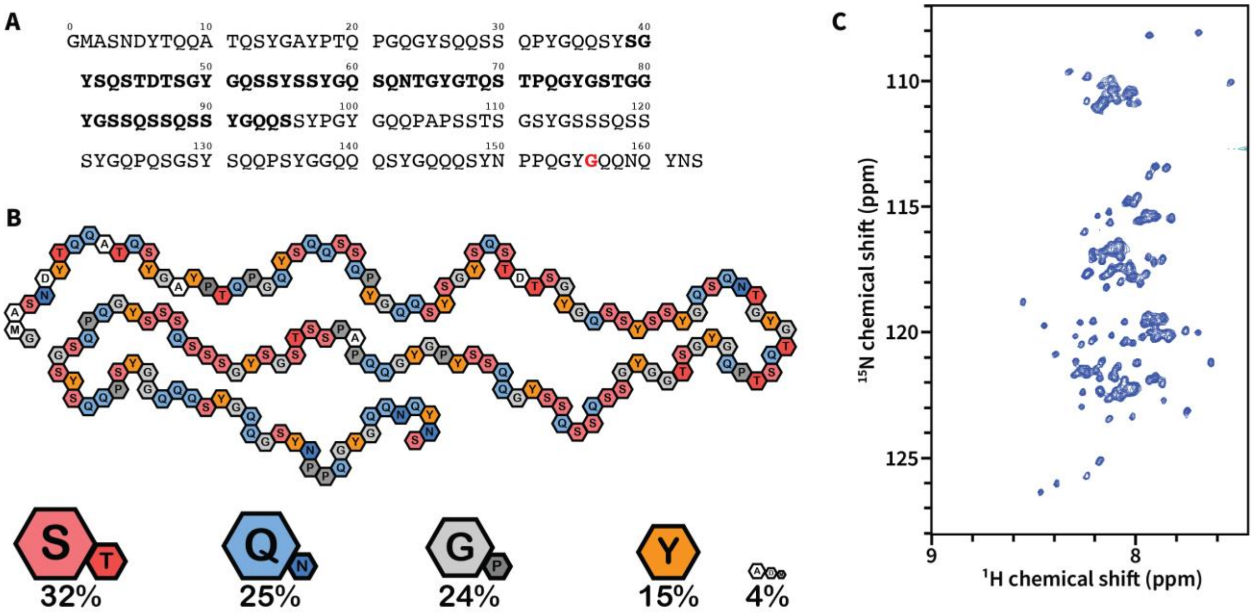


**Figure S1. (A)** The sequence of FUS LC (1 -163) used in this work. The N-terminal glycine is left after cleavage with TEV protease. The seeded fibril core determined by Murray *et al.*[1] is highlighted in bold, and the site of the G156E mutation is marked in red. **(B)** Amino acid composition analysis of the FUS LC construct. **(C)** Solution NMR spectrum of 150 µM FUS LC obtained at 800 MHz in 20 mM MES, pH 5.5 at 25 °C similar to the conditions used by Burke *et al.*[2] where FUS LC can form droplets but not amyloids.

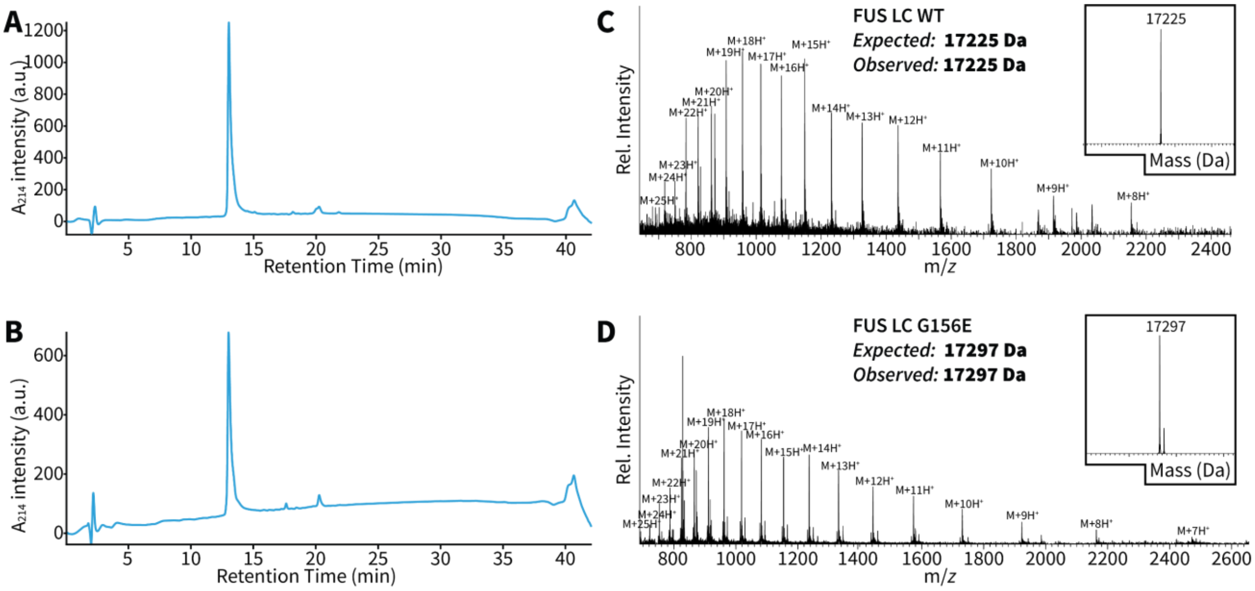


**Figure S2. (A-B)** Representative analytical C18 reverse-phase chromatograms of purified FUS LC and FUS LC G156E respectively. **(C-D)** ESI-MS of the purified proteins.

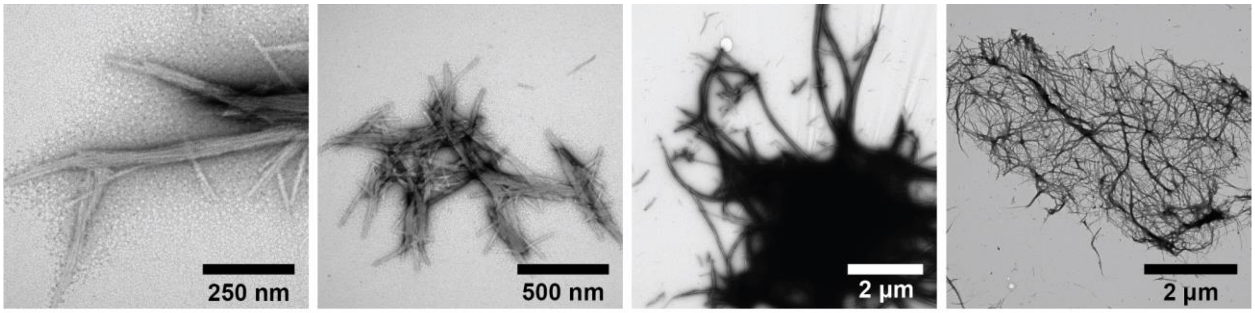


**Figure S3.** Representative transmission electron microscopy (TEM) images of a wild-type FUS LC sample after two months of maturation.

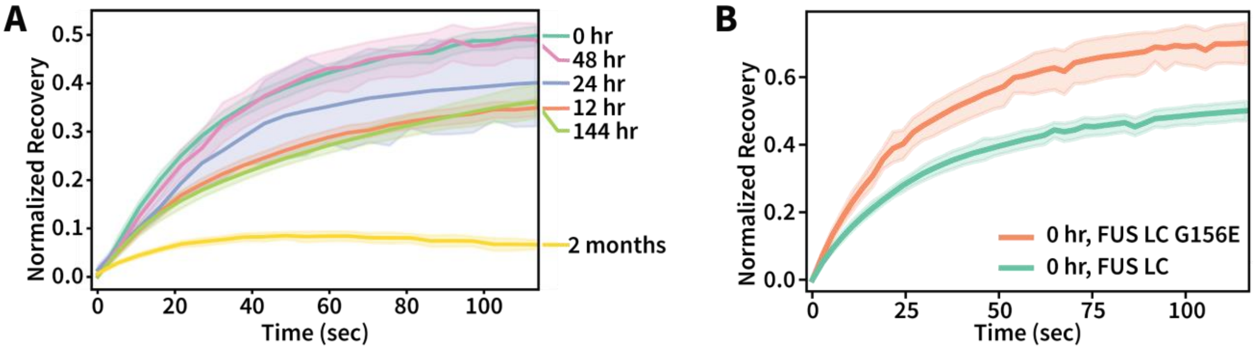


**Figure S4. (A)** FRAP recovery curves of Cy3-labeled FUS LC droplets (5% Cy3-labeled protein) followed over the course of several weeks as they mature into gels and fibrillar species. During the first few days, recovery is highly heterogeneous as not all droplets have reached the same gel-like state. Six unique droplets were photobleached for each time point and data are plotted as the mean recovery with a 95% confidence interval. **(B)** FRAP recovery of FUS LC G156E droplets recorded immediately after LLPS initiation indicating a high degree of mobility at the initial stages of the transformation process. Six unique droplets were photobleached for each time point and data are plotted as the mean recovery with a 95% confidence interval. Recovery time $\tau_{fast}$ = 10 s for the G156E mutant and 20 s for the wild-type sample.

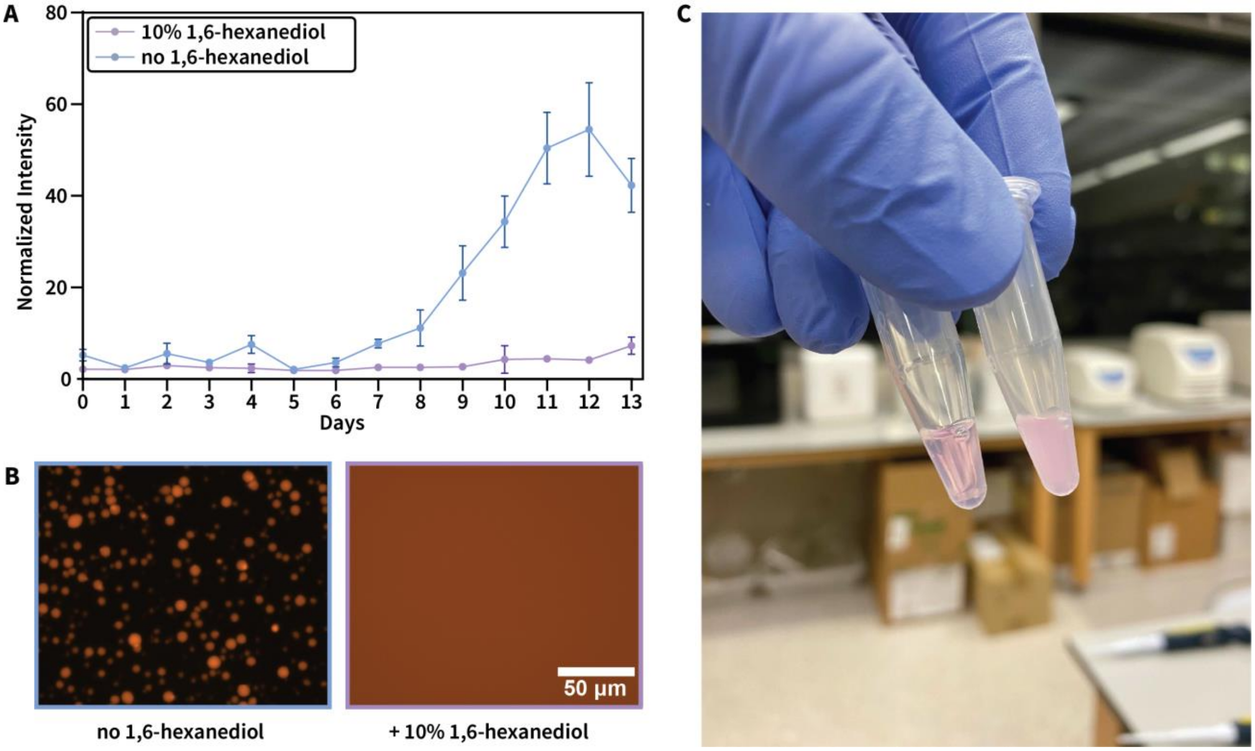


**Figure S5. (A)** Thioflavin T fluorescence assay of FUS LC amyloid formation in the presence and absence of 10% 1,6-hexanediol. Experiments were done in triplicate. **(B)** Imaging of Cy3 labeled FUS LC samples in the presence and absence of 10% 1,6-hexanediol. **(C)** Cy3 labeled FUS LC samples with and without 10% 1,6-hexanediol. Inhibition of LLPS is apparent in the hexanediol sample (left).

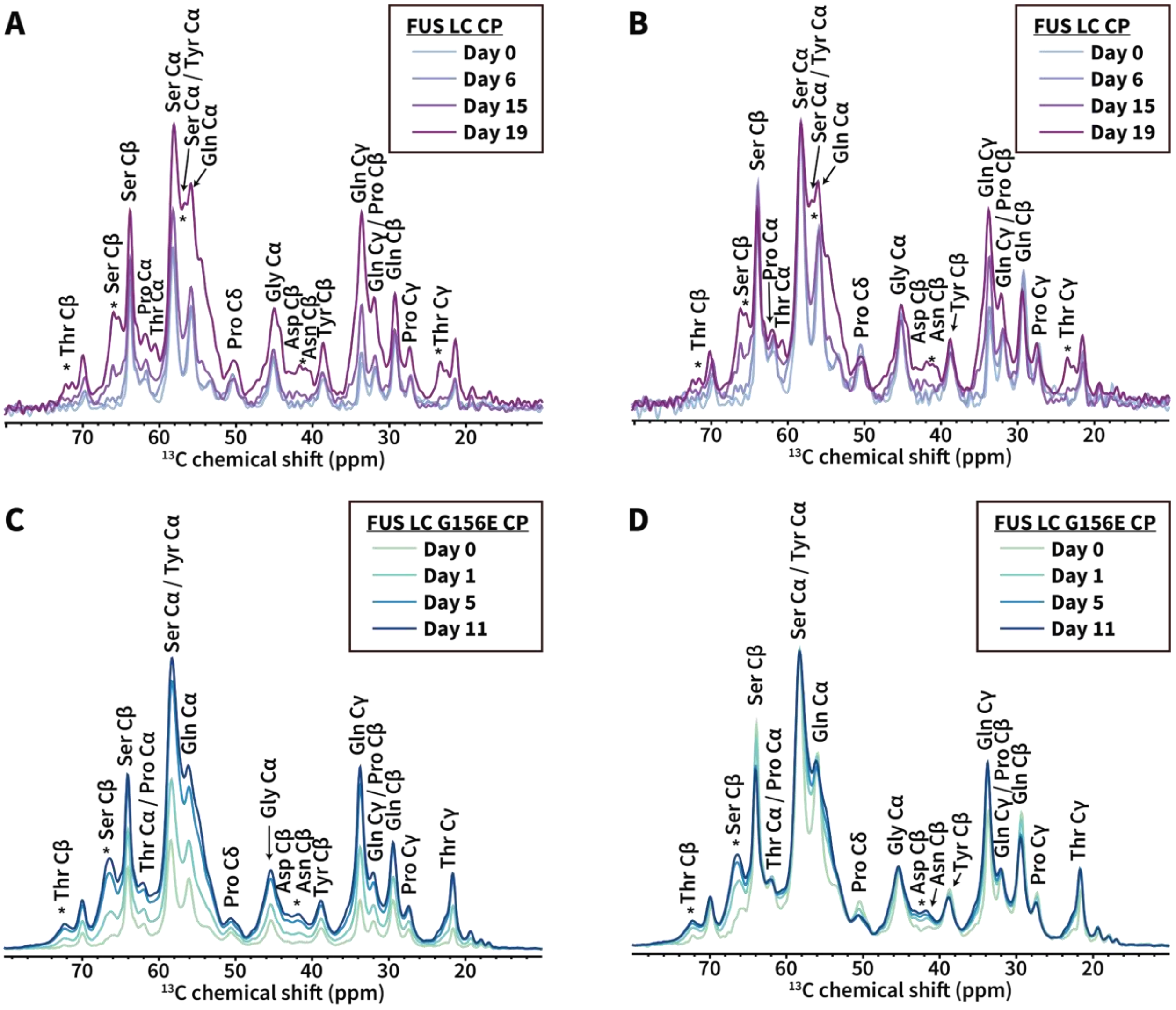


**Figure S6. (A)** Representative CP spectra of the FUS LC sample acquired on different days along the transformation process. **(B)** The same spectra as in (A) but normalized to the highest intensity peak of the day 19 spectrum to highlight changes in chemical shifts throughout the transformation process**. (C)** Representative CP spectra of the FUS LC G165E sample acquired at different days along the transformation process. **(D)** The same spectra as in (C) but normalized to the highest intensity peak of the day 11 spectrum to highlight changes in chemical shifts throughout the transformation process. The asterisks highlight peaks where significant changes are observed.

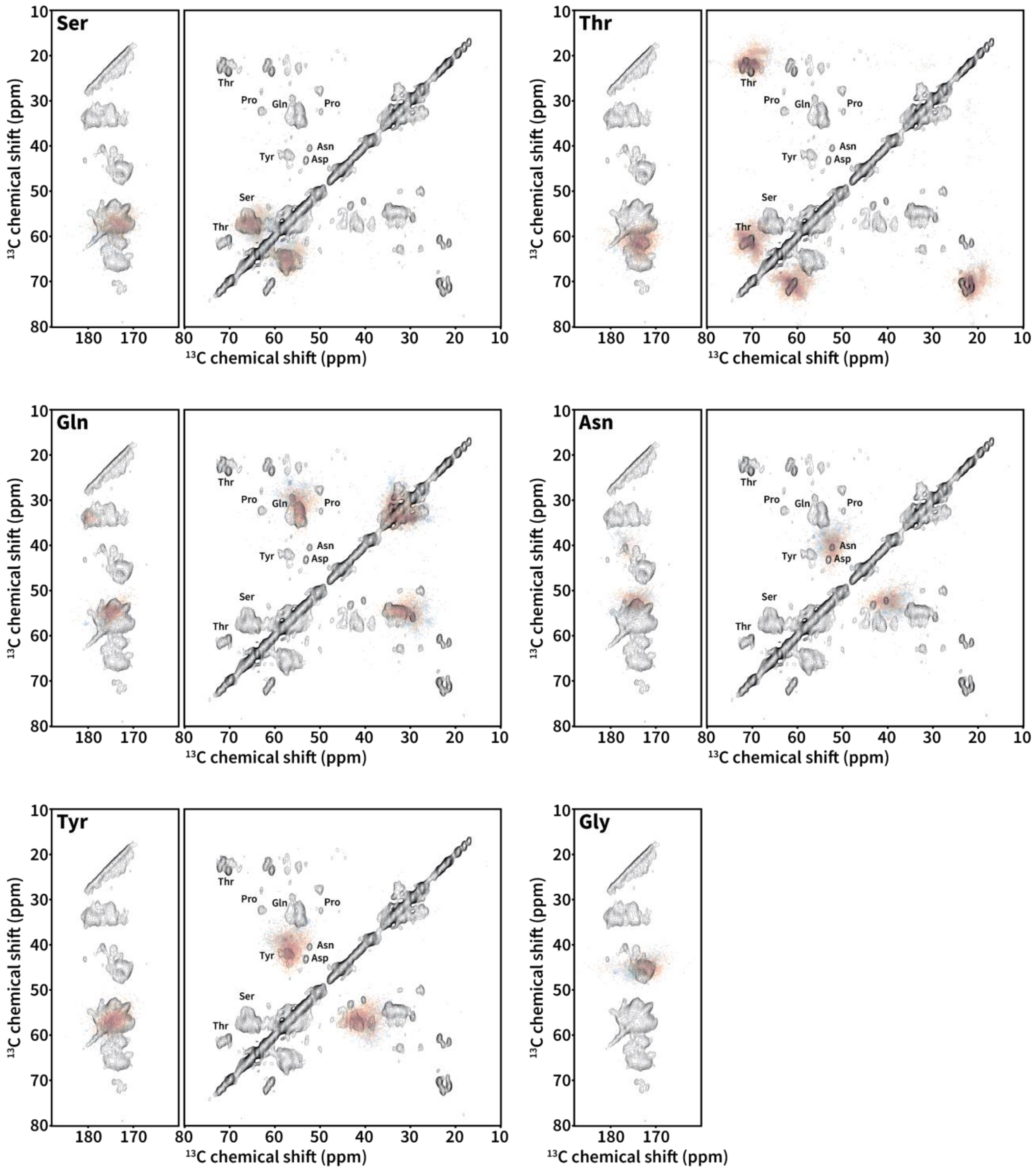


**Figure S7.** Analysis of the β-sheet content of the 2D DARR correlation spectra of FUS LC based on chemical shift statistics from the BMRB. For each residue type, red densities denote the statistical distribution of β-sheet chemical shifts, while blue densities represent random coil chemical shifts. Although there is some overlap between β-sheet and coil densities, the correlations observed in our DARR experiment fall well within the β-sheet statistical distributions, suggesting that the final state of FUS LC contains β-sheet secondary structure.

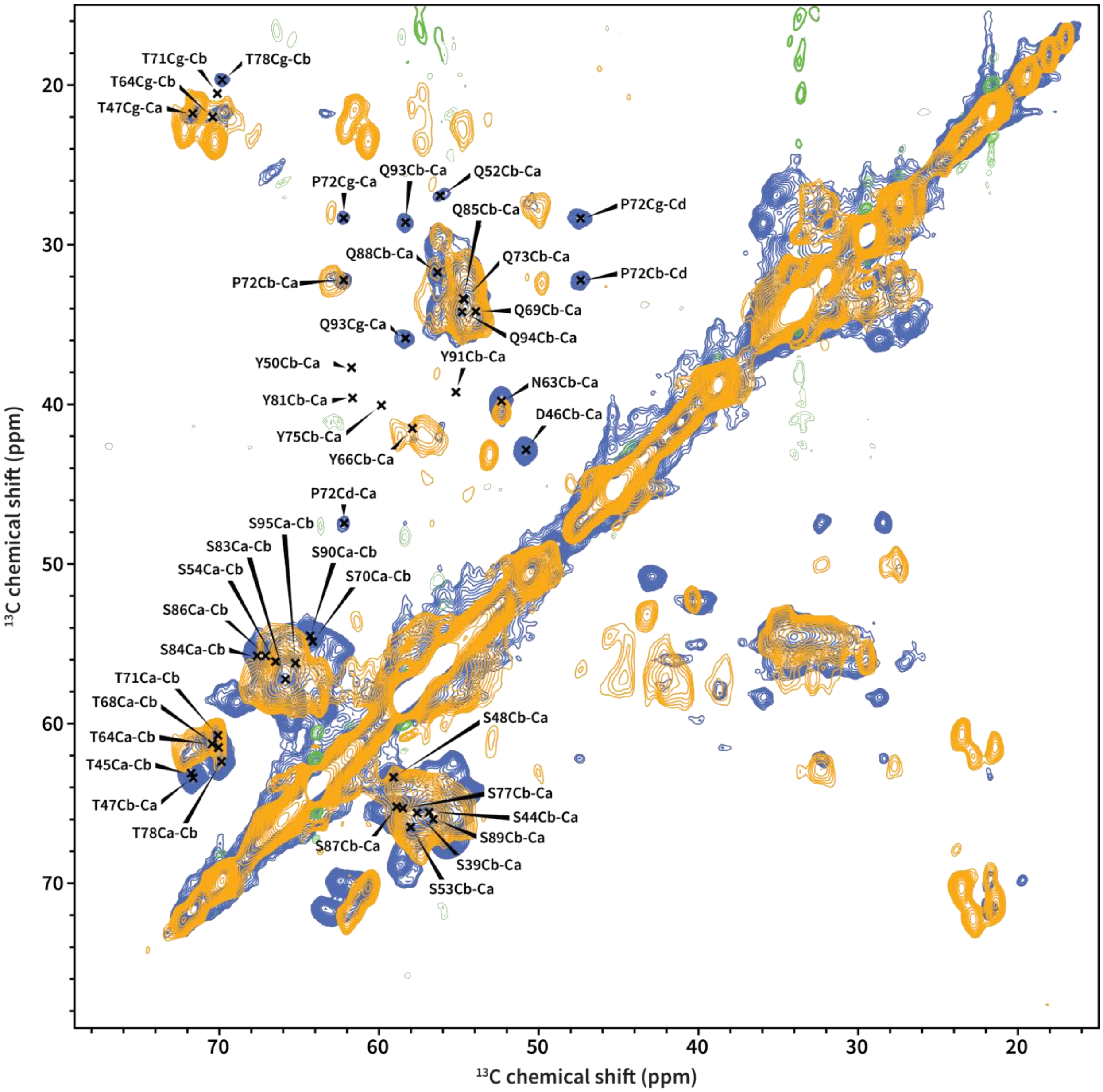


**Figure S8.** Comparison of the DARR spectra of the FUS LC(1-163) sample used in this work (orange) and the DARR spectra of FUS LC(1-214) from Murray *et al*. (blue)[1].The assignments correspond to the FUS LC(1-214) sample as determined by Murray *et al.*

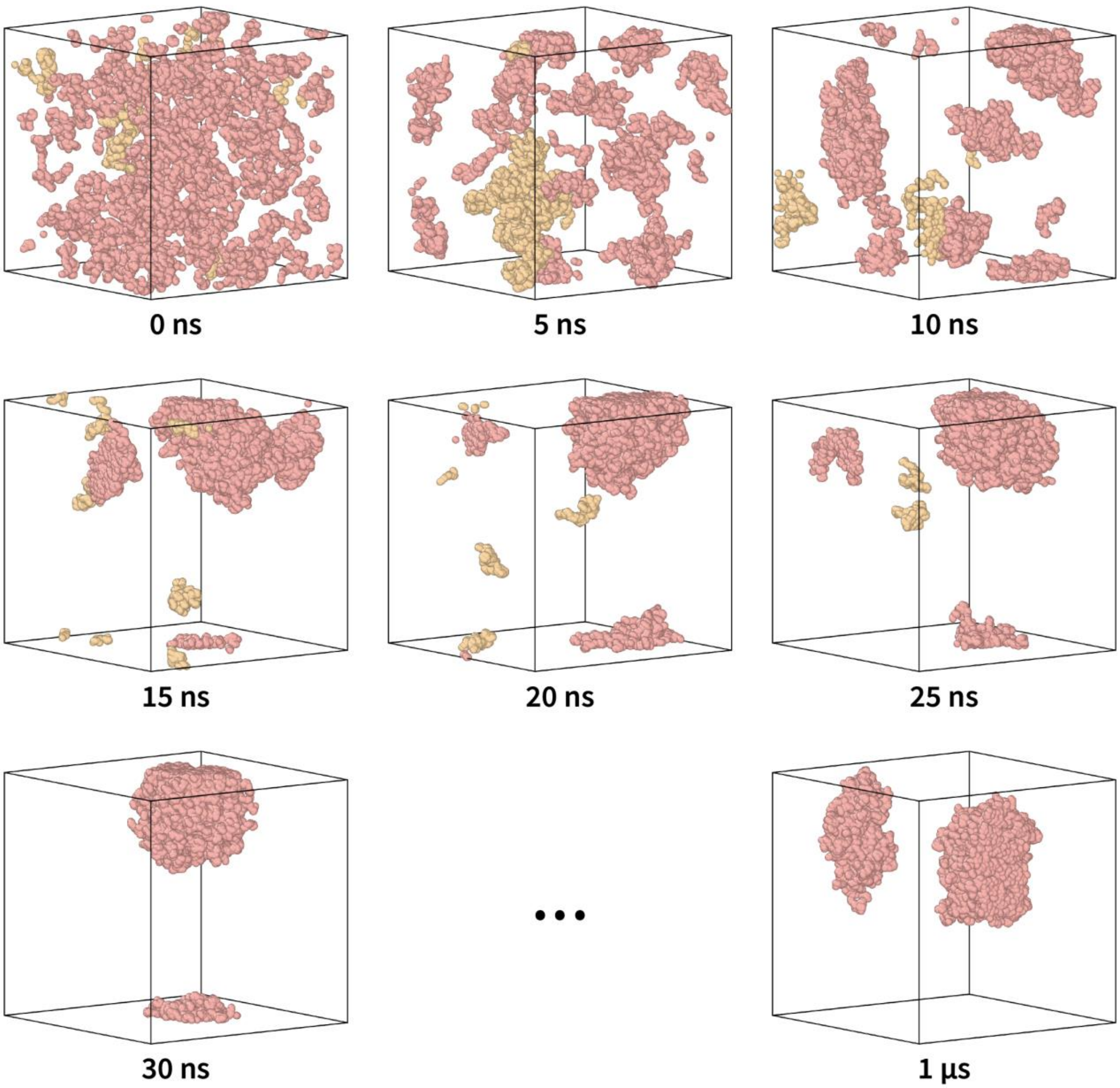


**Figure S9.** Snapshots of the production coarse-grained molecular dynamics run of FUS LC with FUS LC monomers colored by cluster. Clusters are separated by at least 3.2 nm. Particles representing 100 FUS LC monomers were placed into a 50 $nm^3$ simulation cell at the beginning of the run. As the simulation progressed, the protein rapidly condensed into a discrete droplet. The droplet integrity was maintained throughout the entire run, with few monomers occasionally breaking free and then rejoining the droplet.

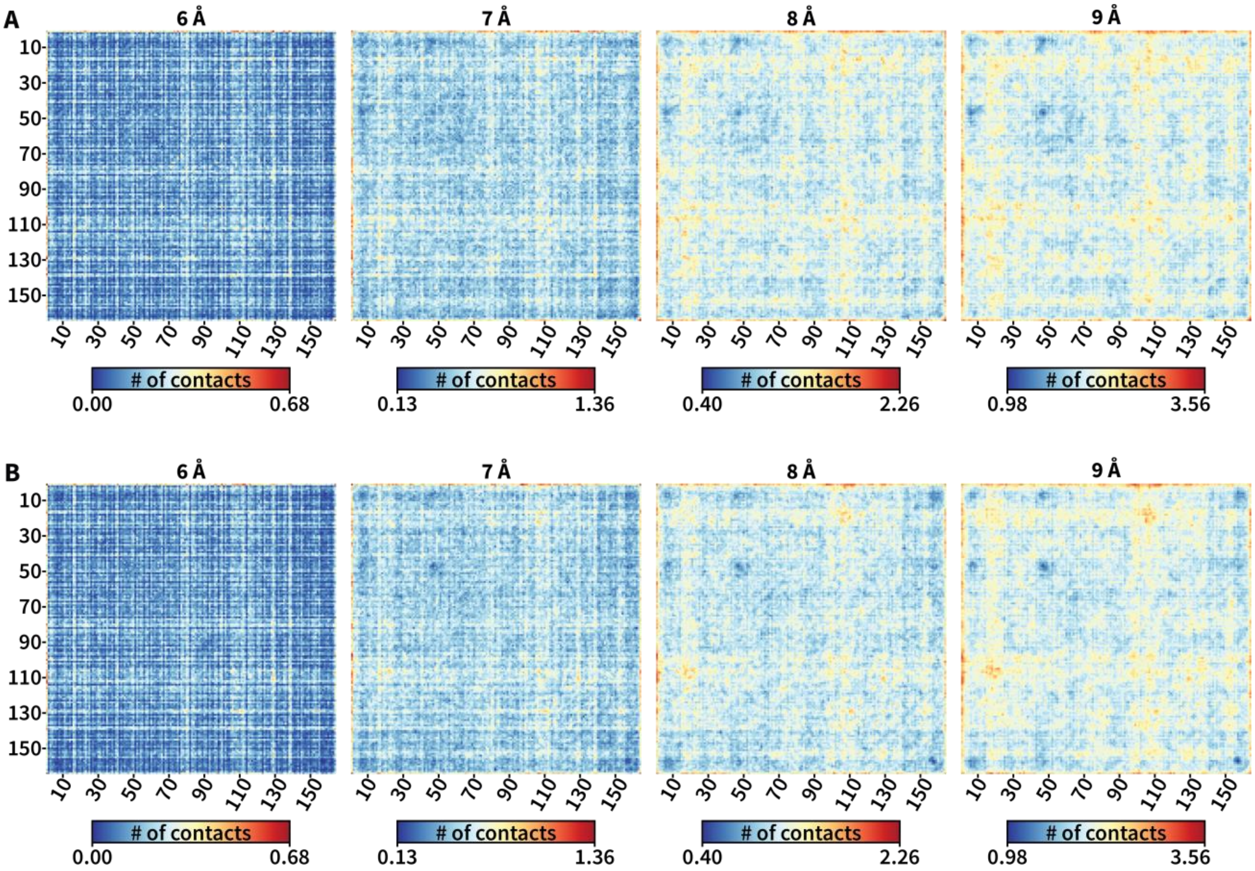


**Figure S10.** Intermolecular contact maps for FUS LC and FUS LC G156E with varying cutoff radii. Distances are measured from bead centers. The contacts maps for each cutoff radius use the same color coding for the number of contacts. **A)** Maps for FUS LC. **B)** Maps for FUS LC G156E.

**Table S1.** Parameters for NMR experiments. All experiments were performed at MAS frequency of 11.11 kHz, 285 K, 750 MHz $^{1}$H Larmor frequency with a 3.2 mm E$^{free}$ HCN probe.

| **Experiment** | **FUS LC** | | **FUS LC G156E** | |
|---|---|---|---|---|
| **Sample amount** | **20** | **mg** | **20** | **mg** |
| CP | | | | |
| Number of scans | 128 | scans | 1600 | scans |
| Recycle delay | 4 | s | 4 | s |
| CP contact time | 1.75 | ms | 1.75 | ms |
| Decoupling | SPINAL64 | | SPINAL64 | |
| Decoupling power | 90 | kHz | 90 | kHz |
| Acquisition time | 9 | ms | 9 | ms |
| INEPT | | | | |
| Number of scans | 128 | scans | 1600 | scans |
| Recycle delay | 4 | s | 4 | s |
| $^{13}$C π/2 | 5 | µs | 5 | µs |
| $^{1}$H π/2 | 2.75 | µs | 2.75 | µs |
| Refocusing parameter | 140 | Hz | 140 | Hz |
| Decoupling | SPINAL64 | | SPINAL64 | |
| Decoupling power | 90 | kHz | 90 | kHz |
| Acquisition time | 9 | ms | 9 | ms |
| DP | | | | |
| Number of scans | 128 | scans | 1600 | scans |
| Recycle delay | 4 | s | 4 | s |
| $^{13}$C π/2 | 5 | µs | 5 | µs |
| Decoupling | SPINAL64 | | SPINAL64 | |
| Decoupling power | 90 | kHz | 90 | kHz |
| Acquisition time | 9 | ms | 9 | ms |
| 2D-INEPT | | | | |
| Number of scans | 32 | scans | 32 | scans |
| Recycle delay | 3 | s | 3 | s |
| $^{13}$C π/2 | 5 | µs | 5 | µs |
| $^{1}$H π/2 | 2.75 | µs | 2.75 | µs |
| Refocusing parameter | 140 | Hz | 140 | Hz |
| Decoupling | swftppm | | swftppm | |
| Decoupling power | 90 | kHz | 90 | kHz |
| t1 acquisition | 33.3 | ms | 33.3 | ms |
| t1 points | 500 | | 500 | |
| CP-DARR | | | | |
| Number of scans | 160 | scans | 64 | scans |
| Recycle delay | 4 | s | 4 | s |
| CP contact time | 1.75 | ms | 1.75 | ms |
| Decoupling | SPINAL64 | | SPINAL64 | |
| Decoupling power | 90 | kHz | 90 | kHz |
| $^{1}$H power during DARR | 11.1 | kHz | 11.1 | kHz |
| DARR mixing | 20 | ms | 20 | ms |
| t1 acquisition | 3.2 | ms | 3.2 | ms |
| t1 points | 256 | | 256 | |

### References

1. Murray, D. T. *et al.* Structure of FUS Protein Fibrils and Its Relevance to Self-Assembly and Phase Separation of Low-Complexity Domains. *Cell* **171**, 615-627.e16 (2017).

2. Burke, K. A., Janke, A. M., Rhine, C. L. & Fawzi, N. L. Residue-by-Residue View of In Vitro FUS Granules that Bind the C-Terminal Domain of RNA Polymerase II. *Mol. Cell* **60**, 231–241 (2015).
